# The Contribution of the membrane-bound complement regulatory proteins CD46 and CD55 in phases of acute lymphocytic leukemia (ALL) and acute myelogenous leukemia (AML)

**DOI:** 10.1101/2024.07.29.601818

**Authors:** Lobna Onsi Saeed, Perihan Ashraf Ammar, Hisham AbdElaziz, Khaled Abou-Aisha, Noha Samir Farag, Mohamed El-Azizi

**Affiliations:** The German University in Cairo; National Cancer Institute, Cairo University

**Keywords:** Acute leukemia, cell viability, CD46, CD55, complement regulatory proteins, flow cytometry, HSB-2 cell line, peripheral blood samples, qRT-PCR, the complement system

## Abstract

The complement system is an essential part of the innate immunity where it is involved in the elimination of pathogens and disposal of apoptotic bodies and immune complexes from the body. Membrane-bound complement regulatory proteins (mCRPs) play key roles in controlling complement activity to avoid any accidental damage of host cells. The role of the complement system during the neoplastic transformation of cells is complicated and has been debated for long. On one hand, the complement system generally acts as a participant in the body’s immune surveillance against cancer. However, recent findings have shown that cancerous cells can use the complement components to assist in certain hallmarks which are fundamental for tumor progression such as angiogenesis, proliferation and metastasis. The aim of the current study is to investigate the differential expression of mCRPs; CD46 and CD55 in of acute lymphoid leukemia (ALL) and acute myeloid leukemia (AML). Clinical peripheral blood samples of newly diagnosed AML and ALL patients were used to assess the changes in transcriptional expression levels of CD46 and CD55 compared to age-matched healthy control subjects using quantitative real time PCR (qPCR). Results showed that both CD46 and CD55 were significantly downregulated by 2 to 7 folds in ALL and AML patients compared to healthy controls. The determined downregulation is suggestive of a defense mechanism conducted by leukemic cells to overcome immune defenses. Flow cytometric analysis was conducted for proteomic expression analysis of both proteins on cell surfaces of leukemia patients compared to healthy controls and results showed a reduction in CD46 expression by 1.2 fold and 2.8 fold in CD55 expression in AML patients. Post transcriptional knockdown of both genes was then carried out in HSB-2 leukemic cell model using customized shRNA, followed by flow cytometry analysis to assess the success of CD46 and CD55 silencing step, and ending with cell viability assays. MTT assay results showed a significant reduction in the viability of HSB-2 cells by 3 fold, approximately, following post-transcriptional silencing of CD46 and CD55 suggesting that although the expression of these mCRPs could be normally compromised by cancerous cells to evade complement attack mechanisms, CD46 and CD55 could be vital to the viability and proliferation of cancerous cells at some point in time. Our results suggest the dual role of complement in the tumor microenvironment where a balance between antitumor and tumor-promoting complement activities exists. The multifunctional properties of the complement system could be employed in opposing roles in cancer suggesting that the biological functions of the complement system are much more diverse than a simple elimination of target cells.

## Introduction

The complement system is a fundamental component of the humoral innate immunity. It helps in elimination of evading pathogens by amplifying inflammatory responses, opsonizing pathogens and attracting phagocytes to the site of infection ^[1][2]^. It is essential for the clearance of immune complexes and apoptotic bodies as well ^[3]^. The complement system components are activated by three major pathways; classical pathway, mannose-binding lectin pathway, and alternative pathway. Complement activation cascade is controlled by complement regulatory proteins (CRPs) in order to protect the host from excessive inflammatory reactions that could lead to host tissue destruction and also to prevent the over-consumption of the complement components ^[1][2]^. CRPs are found in biological fluids or on autologous membranes. Fluid-phase regulators include C1-inhibitor (C1-INH), C4 binding protein (C4bp), Factor I (fI) and Factor H (fH). Membrane-bound complement regulatory proteins (mCRPs) include: Complement Receptor 1(CR1), Membrane co-factor protein (MCP), also known as CD46, Decay Accelerating Factor (DAF), known as CD55, and protectin (CD59)^[4][1]^.

CD46 and CD55 control the complement activation cascade on the level of C3 and C5 convertases ^[5]^. CD46 is widely distributed on human peripheral blood cells except erythrocytes, as well as being expressed on fibroblasts, epithelial and endothelial cells ^[6]^. It controls complement activation by acting as an intrinsic cofactor for Factor-I-mediated cleavage of C3b and C4b ^[7][8]^. Cleavage of C3b by Factor I in presence of CD46 produces iC3b, which cannot support any further complement activation. While cleavage of C4b produces C4c which is released and C4d which remains bound on the cell surface with no activity ^[9]^. CD46 also plays a role in cellular-mediated immunity where it helps in the differentiation of CD4^+^ T lymphocytes into regulatory T cells producing IL-10 that suppresses T helper cells ^[10]^.

As for decay accelerating factor (DAF); also known as CD55, it is a type I cell surface protein that forms a single chain anchored to the membrane by glycosylphosphatidylinositol (GPI). It binds C3b and C4b inhibiting thereby the formation of C3 convertase and decreasing its half-life, thus providing a protective barrier threshold for plasma membranes of normal autologous cells against complement deposition and activation ^[11][12]^.

The role of the complement system in cancer is complicated and has been debated for long. Malignant transformation is generally accompanied by genetic and epigenetic modifications which drastically alter patterns of glycosylation, cell-surface proteins and phospholipids ^[13][14]^. These alterations can be identified by innate and adaptive immune mechanisms that guard the host against cancer development ^[15]^. This is the known basis of the immune surveillance hypothesis. There is no direct evidence to support the argument that complement is able to eradicate emerging tumors. Nevertheless, taking into consideration that complement is intended for the recognition of non-self-elements, it is assumed that alterations in the tumor cell membranes’ composition render these cells as targets for complement recognition ^[16]^. However, the relationship between inflammation and cancer is complicated and subject to contradictory forces ^[17]^. Therefore, while acute responses are considered a vital part of the defense against cancerous cells, continuous inflammation in the tumor microenvironment increases the threat of neoplastic transformation and has several tumor-promoting effects ^[18][19]^.

The current study aims at investigating the expression levels of mCRPs; CD46 and CD55 in the acute lymphocytic leukemia and acute myelogenous leukemia and to further elucidate its role in Egyptian cancer patients. To the best of our knowledge this is the first study conducted on Egyptian leukemia patients and one of very few studies tackling the complicated role of complement in acute leukemia.

## Materials and Methods

### Study group

Thirty peripheral blood samples were collected from acute lymphocytic leukemia (ALL) and acute myeloid leukemia (AML) patients of the National Cancer Institute, Cairo University. 14 samples were taken from patients with ALL and 16 samples from patients with AML. The age range was adults (20 - 63 years old) of both genders. All cases were newly diagnosed, have taken no or minimal chemotherapy and with a minimum of 40% leukemic blasts present in peripheral blood. Eight control blood samples were also collected from healthy volunteers within the same age range. The study was approved by the Human Ethics Review Committee of the Faculty of Medicine, Cairo University and participants were informed in detail about the study and gave their written consents. Participation in the study was voluntary and informed.

### Total RNA extraction from blood samples and reverse transcription of total RNA into complementary DNA (cDNA)

Samples were collected in EDTA containing vacutainers with RNAlater (ThermoFischer Scientific, USA). Total RNA was extracted using Trizol reagent (ThermoFischer Scientific, USA) following Genomic Medicine Biorepository (GMB) protocol^[20]^. Each RNA pellet was then dissolved in 20µl of RNAse-free water and stored at −80°C. RNA concentration was determined fluorometrically using Qubit 4 (ThermoFisher Scientific, USA) and its integrity was tested using 1% agarose gel electrophoresis.

Reverse transcription was then performed using High Capacity cDNA Reverse transcription Kit (Applied Biosystems, USA) following the kit’s manual ^[21]^, using 2 µg of the total RNA.

### Assessing gene expression of CD46 and CD55 in acute leukemia patients by quantitative real-time polymerase chain reaction (qRT-PCR)

Gene expression analysis for CD46 and CD55 was done using Taqman gene expression assays Hs00611257_m1 and Hs00892618_m1 (Thermofisher scientific, USA), respectively, with Beta Actin (Hs01060665 g1) being the housekeeping or reference gene. Results were interpreted using the comparative ΔΔCt method^[22]^.

### Flow cytometric analysis of CD46 and CD55 proteins expression levels on cell surfaces of leukemia clinical samples

The expression levels of CD46 and CD55 proteins in AML and ALL patient blood samples were analyzed using FACS assay monoclonal antibodies; CD46 labelled with FITC and CD55 labelled with PE (A15745 and 12055942, Thermo Fisher Scientific). Briefly, blood cells (1 × 10^5^) were pelleted at 2000 rpm, supernatant was discarded, and pellet was washed in 100 mL FACS-buffer (1% BSA, 0.1% NaN3 in PBS) and centrifuged at 2000 rpm, for 4 minutes. Then supernatant was removed followed by addition of the corresponding monoclonal antibody. The cells were washed twice with FACS buffer for removal of any excess dye and pellet was finally resuspended in PBS buffer. Stained cells were analysed by FACS Calibur (BD Biosciences, USA) using Cell-Quest Pro Software Results were compared with the expression levels of the targeted proteins in healthy controls.

### Post-transcriptional knockdown of CD46 and CD55 in HSB-2 leukemic cell line using shRNA

HSB2-ALL cell line was cultured in Roswell Park Memorial Institute medium (RPMI) + L-glutamine medium supplemented with 10% fetal bovine serum and Pen-Strep and maintained at 37°C, 5% CO_2_. Cells density was kept between 2×10^5^ cells/ml and 1×10^6^ cells/ml.

shRNA plasmids purchased from Santa Cruz Biotechnology, Inc. were used to knockdown the expression of CD46 and CD55 (Tables 1 and 2, respectively). Moreover, Sc-108060 shRNA plasmid-A (encoding for a scrambled sequence) was used as a mock control to eliminate any nonspecific effects that may be caused by the transfection reagent or process. Each plasmid consisted of a pool of 3 different shRNA plasmids to ensure high transfection efficiency (Tables 1 and 2).

**Table 1.**
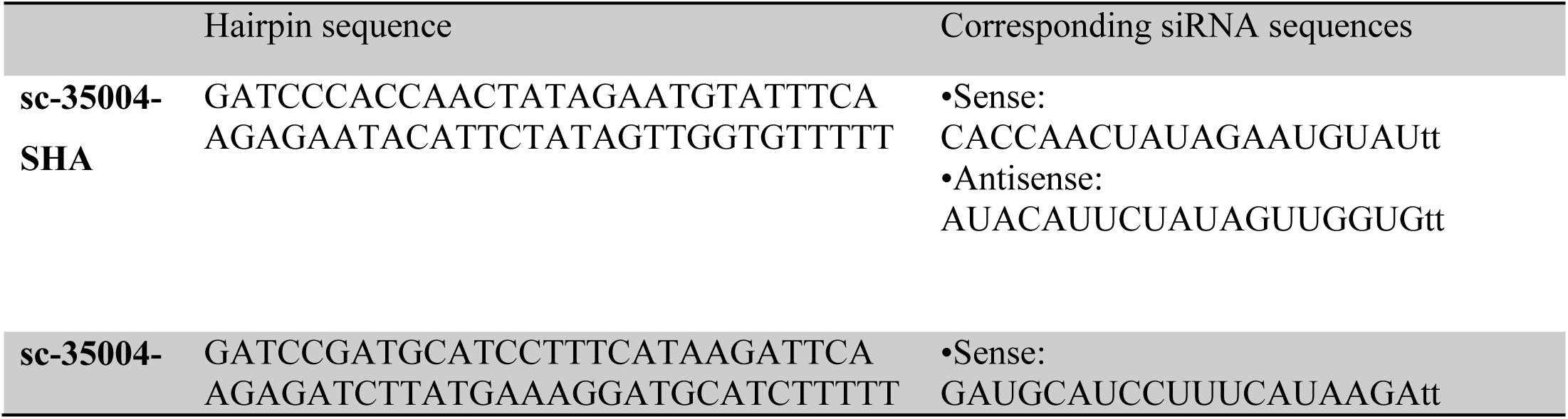
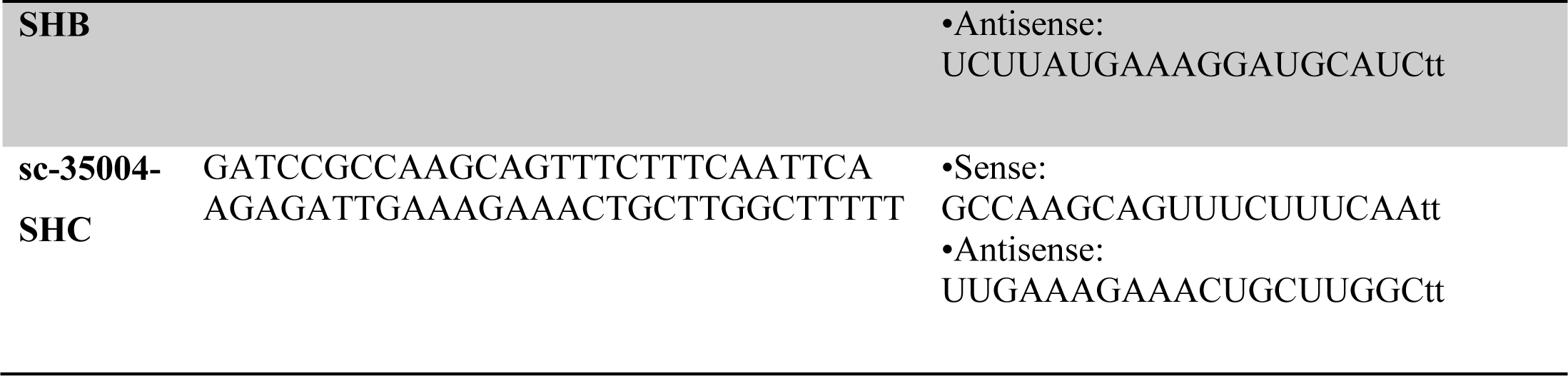
CD46 shRNA (sc-35004-SH) Plasmid (h) is a pool of 3 different shRNA plasmids.

**Table 2.**
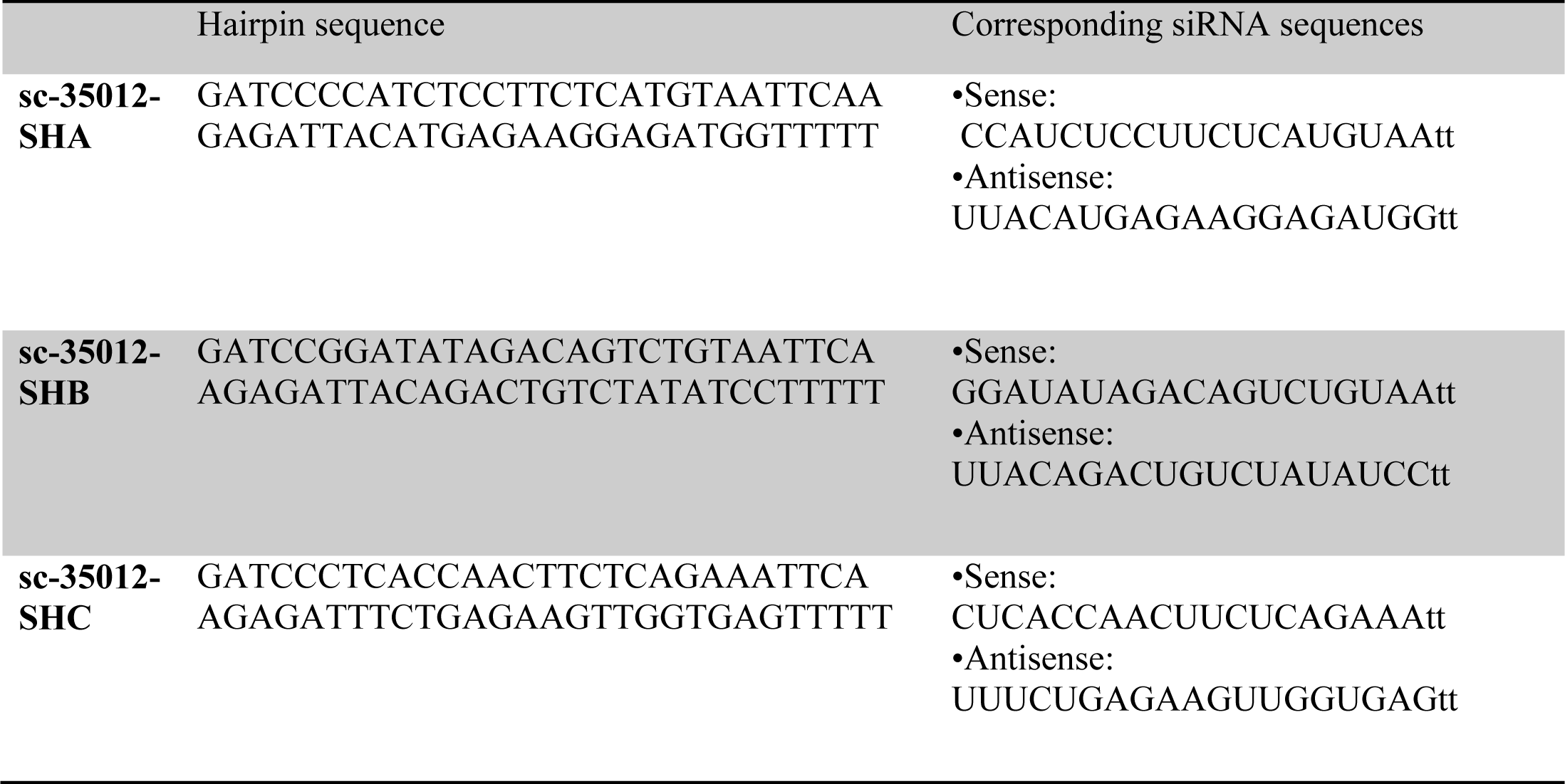
CD55 shRNA (sc-35012-SH) Plasmid (h) is a pool of 3 different shRNA plasmids.

HSB-2 cells were transfected using polyethylenimine (PEI) reagent. Cells were seeded in a 6-well plate before transfection until 70%-80% confluency. Two solutions were prepared for transfection; solution A consisting of the shRNA plasmid of choice with a concentration of 0.1µg/µl in RPMI in a ratio of 1:3 and solution B consisting of PEI in RPMI in a ratio of 1:1. 1µg/ml of puromycin antibiotic was added for selection. For combined transfection, both CD46 and CD55 shRNA plasmids were added together to observe the effect of co-silencing on cell viability.

### Flow cytometry analysis of post-transfection expression levels of CD46 and CD55

Cells were washed in phosphate buffer saline. CD46 labelled with FITC and CD55 labelled with PE monoclonal antibodies (ThermoFisher scientific, USA) were used. Antibodies were added to the cells in a ratio of 1:10 and they were incubated for 15 minutes. Afterwards, they were washed with PBS to remove excess fluorescent dyes before performing flow cytometry.

### Cell viability assay

Transfected cells seeded in a 6-well plate were incubated for 24 hours in a complete culture medium containing puromycin as well as normal human serum (NHS) freshly prepared from healthy blood donors as a source of complement. Cells were later seeded in a 96-well plate. MTT was prepared in PBS at a concentration of 5 mg/ml and filter sterilized. Cells were cultured for three days at 37°C, 5% CO_2_ and then MTT assay was performed 3 days after culture. To each well 100 µl of MTT was added and then incubated for 4 hours at 37°C, 5% CO_2_. At the end of incubation, MTT was discarded and100 µl DMSO was added to each well. Cell mixture was resuspended repeatedly to dissolve the precipitate. The plate was then transferred to an ELISA plate reader (Victor3, Perkin Elmer, USA) and absorbance was measured at 595nm.

### Statistical analysis

Using GraphPad Prism software (version 8), the data in this study are presented as the mean ± standard deviation and analysis of different groups was done by one-way ANOVA. (P<0.05) was considered a significant difference.

## Results

### Assessing gene expression of CD46 and CD55 in acute leukemia patients by quantitative real-time polymerase chain reaction (qRT-PCR)

CD46 expression was found to be significantly lower (p<0.001) in both AML by 76% compared to healthy controls, and ALL patients (81.7%; p<0.001). Similar results were observed for CD55 where gene expression level was significantly down regulated by 75.1% (p<0.001) in both AML and ALL patient groups as compared to healthy controls (Figure 1).

**Figure 1.**
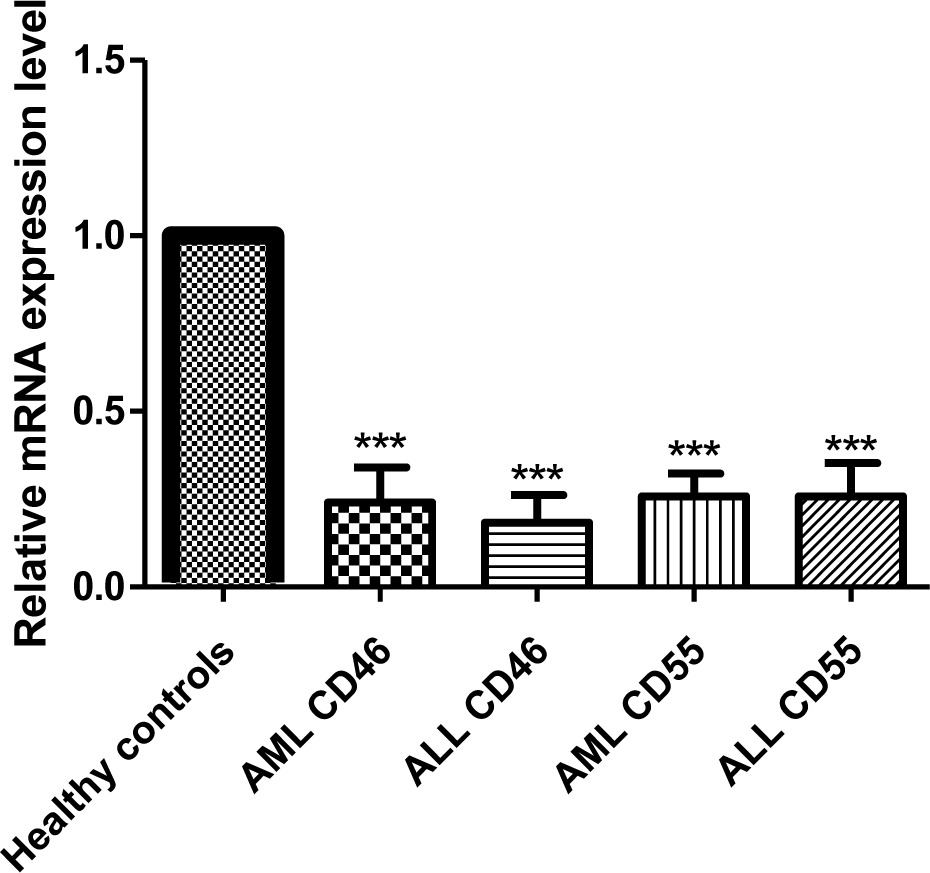
CD46 and CD55 mRNA expression in acute leukemia patients. CD46 and CD55 mRNA expression levels were measured using qRT-PCR in both AML and ALL patients, normalized to the housekeeping gene Beta-actin and results were compared to CD46 and CD55 mRNA expression levels in healthy controls by calculating the RQ value for each one of them. mRNA expression of mCRPs was significantly reduced in AML and ALL patients in comparison with healthy controls. Data are expressed as means ± SD of three independent experiments for each sample done in duplicates and statistical one-way analysis of variance (ANOVA) was used to estimate the significance of difference between the different groups. P<0.001(***).

Furthermore, the level of CD46 and CD55 gene expression was compared in male and female leukemia patients to investigate if gender difference affects their gene expression. Male patients had a slightly reduced expression level of CD46 compared to female patients in both ALL and AML, while for CD55, expression was slightly lower in AML female patients compared to males and on the other hand, expression pattern was reversed in ALL patients (Figure 2).

**Figure 2.**
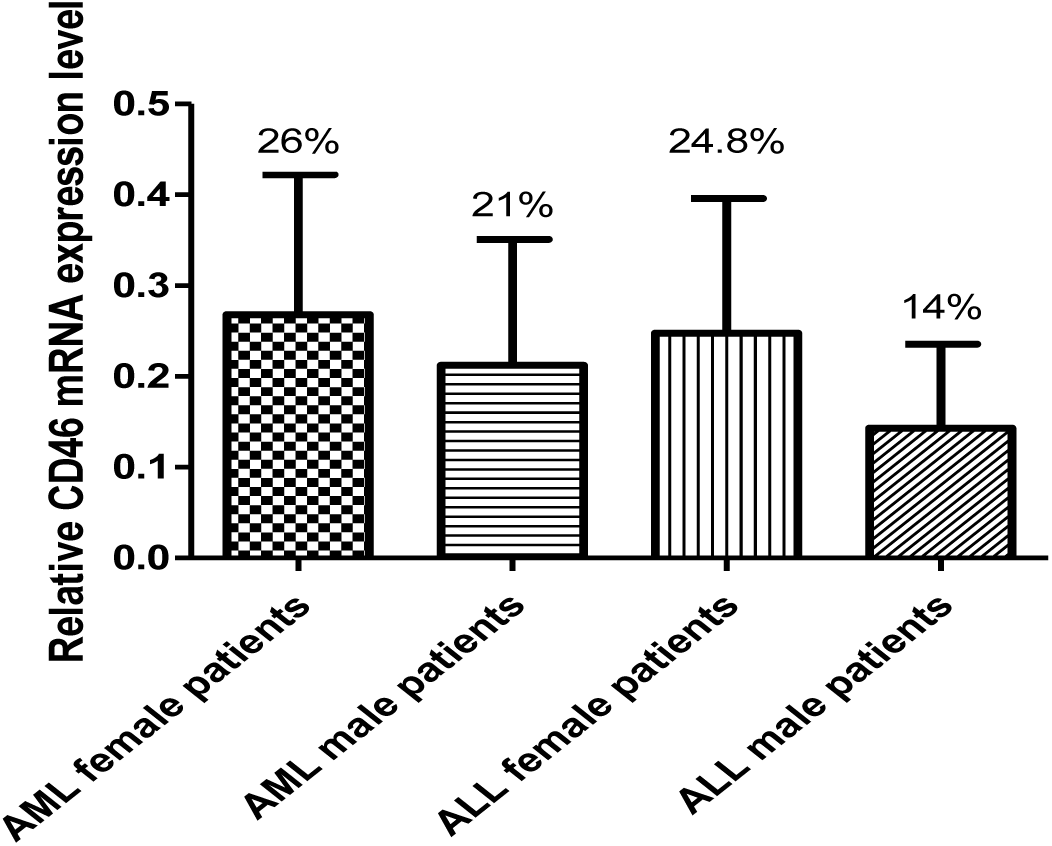
CD46 mRNA expression level based on gender difference. CD46 expression levels compared in AML and ALL male and female patients showed that male patients have a reduced expression level of CD46 compared to female patients in both AML and ALL. Data are expressed as means ± SD of three independent experiments for each sample done in duplicates and statistical one-way analysis of variance (ANOVA) was used to estimate the significance of difference between the different groups.

There was no significant difference (p>0.05) in the expression of CD55 gene between male and female subjects in both AML and ALL groups (Figure 3).

**Figure 3.**
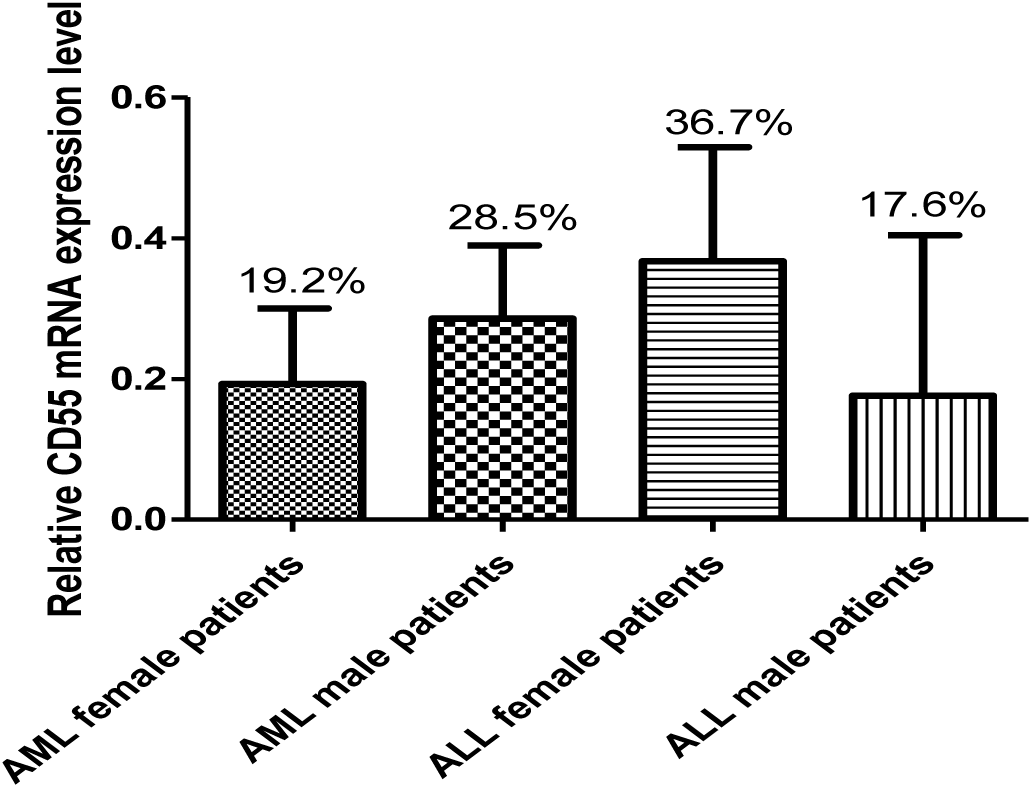
CD55 mRNA expression level based on gender differences. The expression level of CD55 in males and females showed a slight reduction in female AML patients compared to males, while the opposite was observed in ALL patients. Data are expressed as means ± SD of three independent experiments for each sample done in duplicates and statistical one-way analysis of variance (ANOVA) was used to estimate the significance of difference between the different groups.

### FACS analysis of CD46 and CD55 proteins expression levels in peripheral blood samples

FACS analysis confirmed a significant reduction in CD46 protein expression level compared to healthy controls (p<0.05), where the protein expression values were 82.8% and 80.65% for AML and ALL, respectively. As for CD55, a significant reduction was observed in the protein expression level in AML patients compared to healthy controls (expression value of 35.95%) while no significant difference in protein expression was observed between ALL patients and healthy controls (p>0.05) (Figures 4 and 5).

**Figure 4.**
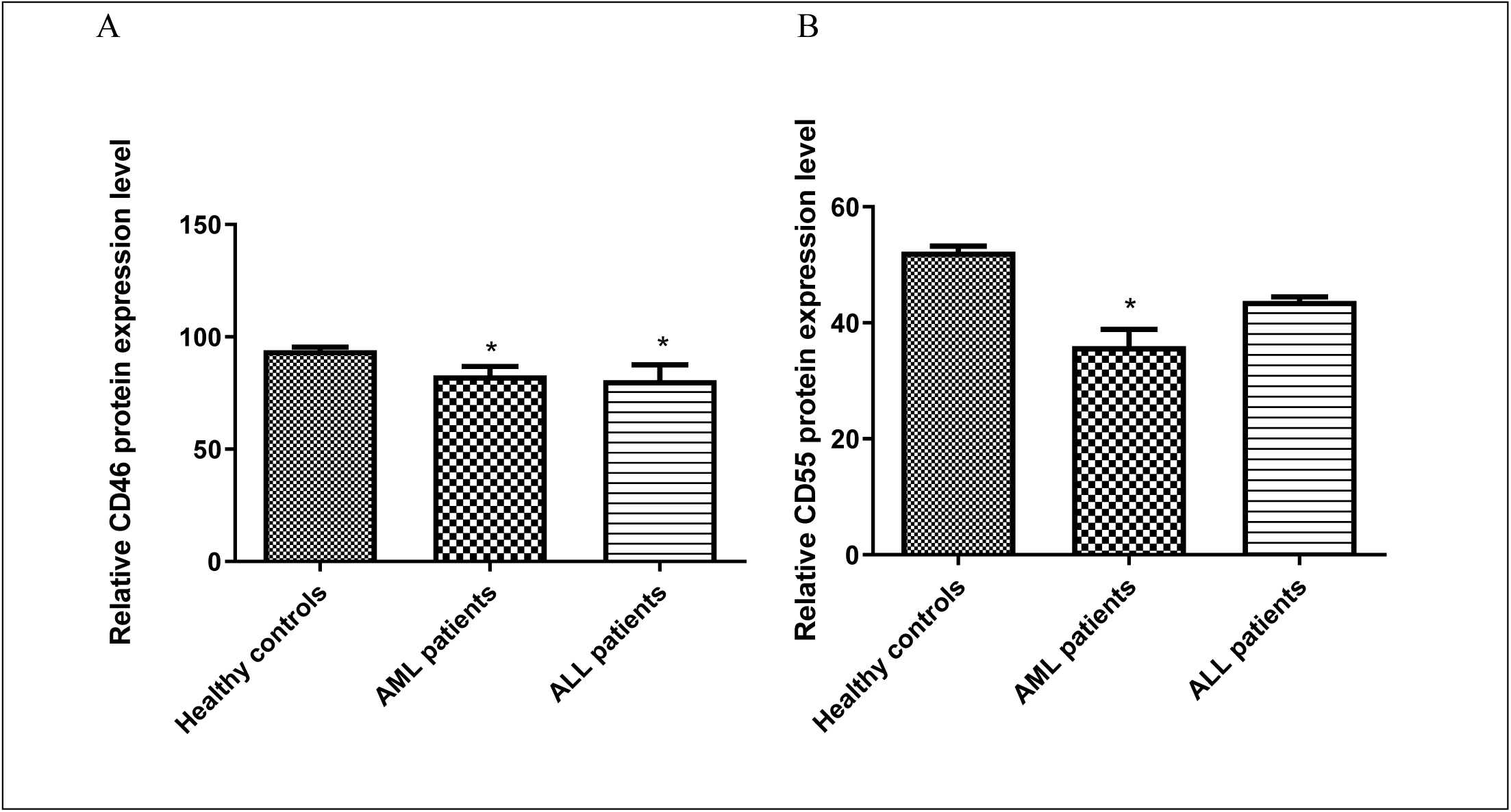
Flow cytometric analysis of the relative expression level of CD46 and CD55 proteins in AML and ALL patients. Flow cytometric analysis performed to measure the expression level of CD46 and CD55 in AML and ALL patients showed a significant reduction in CD46 expression in both types of acute leukemia patients. However, for CD55 a significant reduction was observed in AML patients only, while there was no significant difference observed in case of ALL patients. The expression level was calculated with reference to healthy controls. Data are represented as mean values ±SD for each sample done in duplicates, p<0.05 (*).

**Figure 5.**
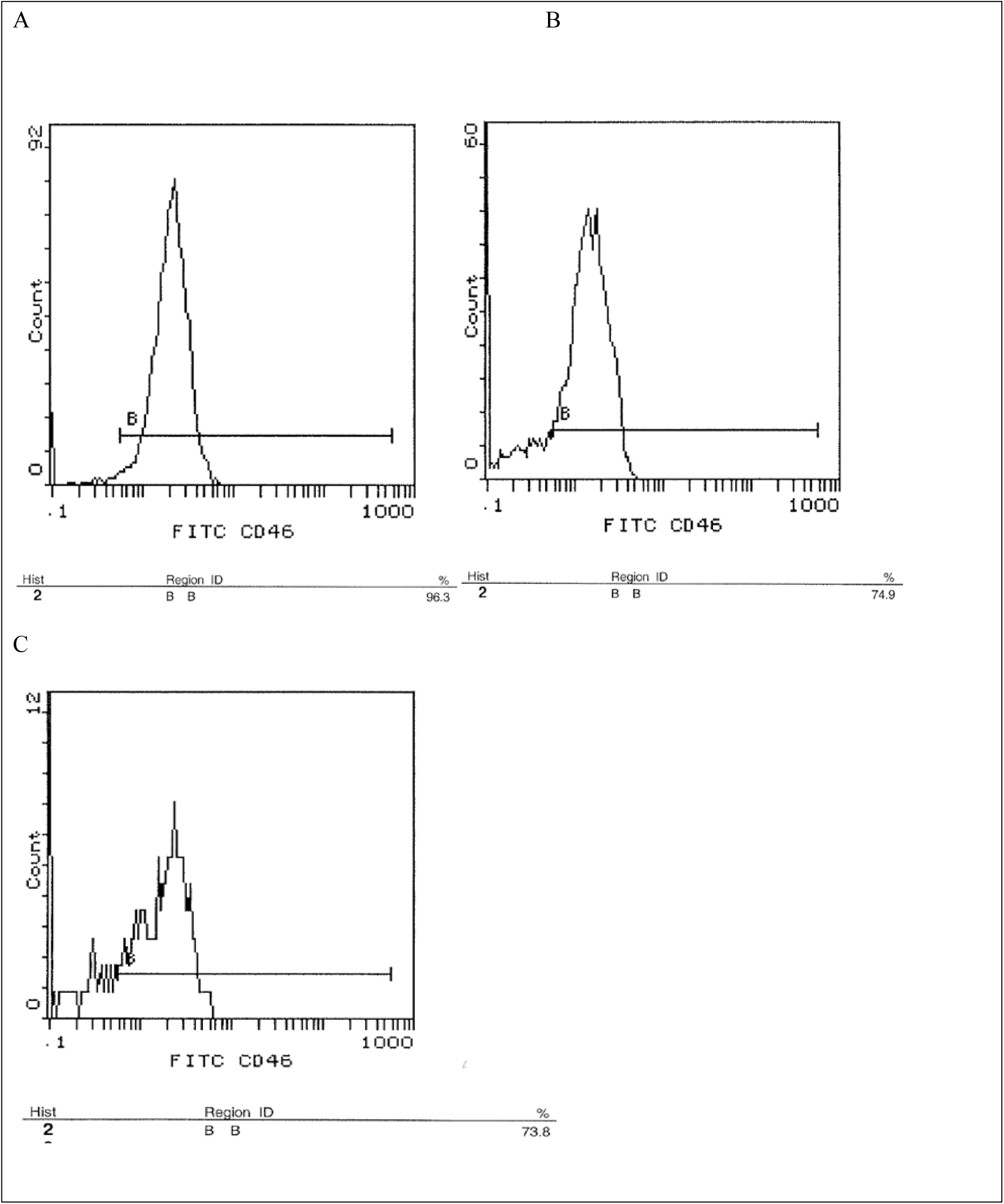
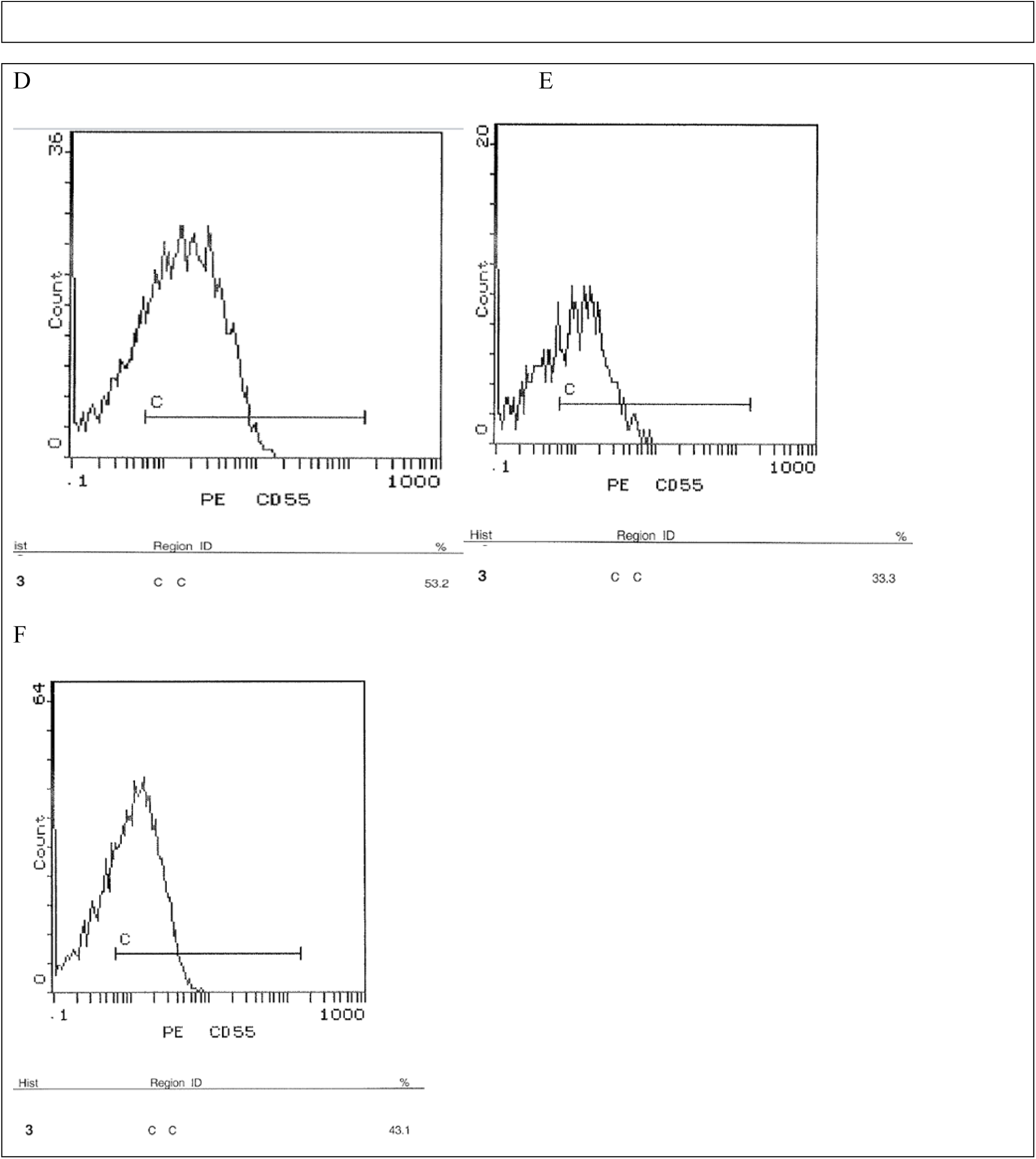
Flow cytometric analysis charts of CD46 and CD55 in healthy controls and acute leukemia patients’ peripheral blood samples. Flow cytometric analysis of the protein expression level of both CD46 and CD55 in the peripheral blood samples of acute leukemia patients showed a significant reduction in both proteins compared to healthy controls. Results shown are from a representative experiment. **(A)** CD46 expression in a healthy control. **(B)** CD46 expression in an AML patient. **(C)** CD46 expression in an ALL patient. **(D)** CD55 expression in a healthy control. **(E)** CD55 expression in an AML patient. **(F)** CD55 expression in an ALL patient.

### ShRNA transfection of HSB-2 leukemic cell line for post transcriptional knockdown of CD46 and CD55

Flow cytometric analysis performed to measure the success of transfection showed a highly significant reduction in CD46 expression level following ShRNA**-**transfection by 91.42% compared to the mock transfected cells. Similar results were observed for CD55 where ShRNA**-** transfection resulted in a significantly reduced CD55 expression level by 87.36% (Figure 6).

**Figure 6.**
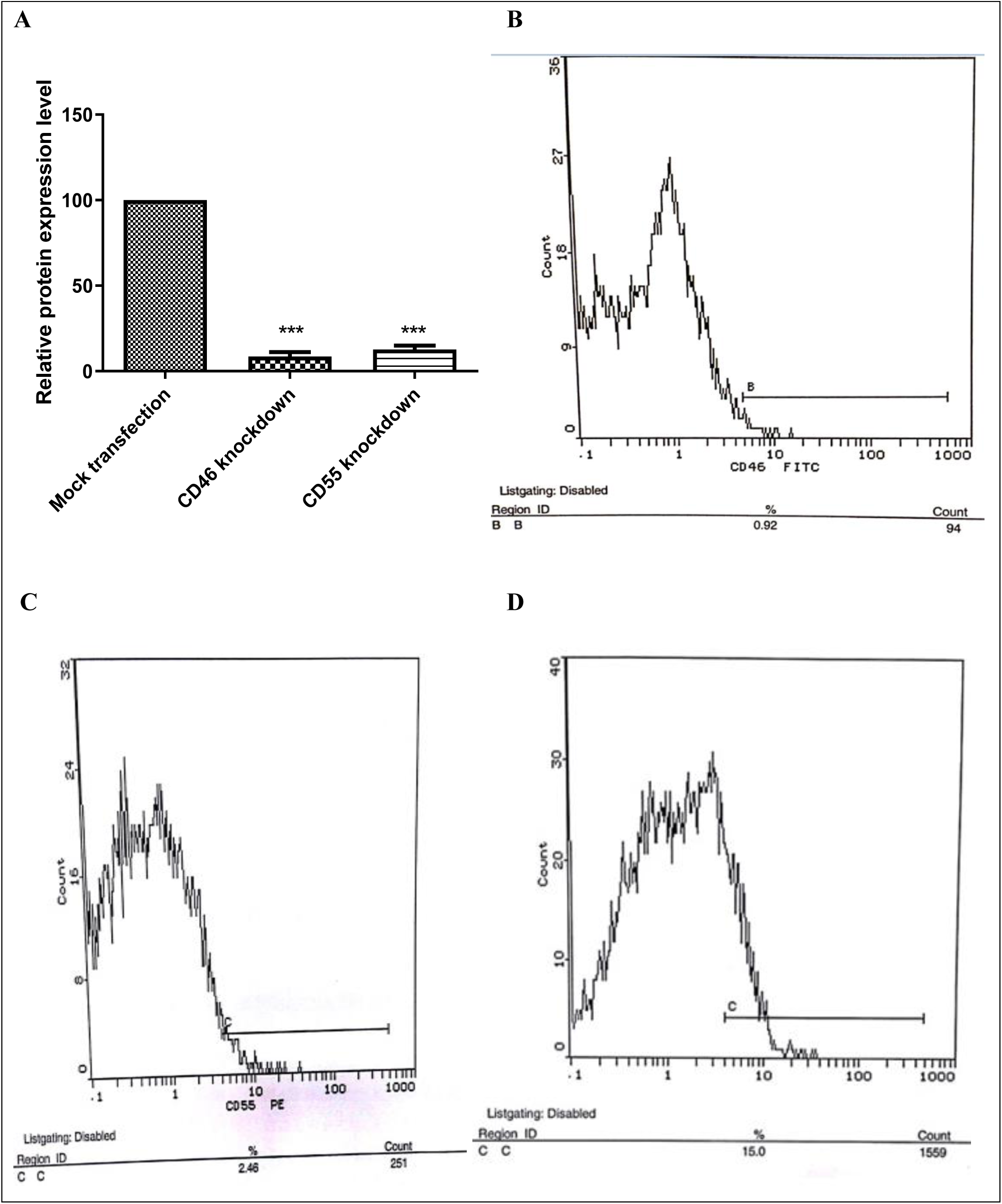
ShRNA-mediated knockdown of CD46 and CD55 mCRPs expression on HSB-2 cells compared to the mock transfected controls. Knockdown of CD46 and CD55 protein expression in transfected cells was performed using shRNA silencing plasmids, each consisting of a pool of three different shRNA plasmids and PEI was used as the transfection reagent. Silencing of CD46 and CD55 proteins was done separately in addition to co-silencing of both proteins. Flow cytometric analysis was performed to measure the expression level of both proteins post transfection. Results showed a highly significant reduction in the expression level of both proteins. The percentage of inhibition was calculated relative to nonsilencing shRNA controls (=100%) p<0.001(***). **(A)** CD46 and CD55 proteins expression level post transfection compared to expression in mock transfected cells. **(B)** Flow cytometric analysis chart of a representative experiment showing protein expression of CD46 in HSB-2 transfected cells. **(C)** Flow cytometric analysis chart of a representative experiment showing protein expression of CD55 in HSB-2 transfected cells. **(D)** Flow cytometric analysis chart of a representative experiment showing protein expression of CD46 and CD55 in mock transfected cells.

Combined knockdown of both CD46 and CD55 resulted in silencing of CD46 by 95.6% (p<0.001) and CD55 by 83.36% (p<0.001) (Figure 7).

**Figure 7.**
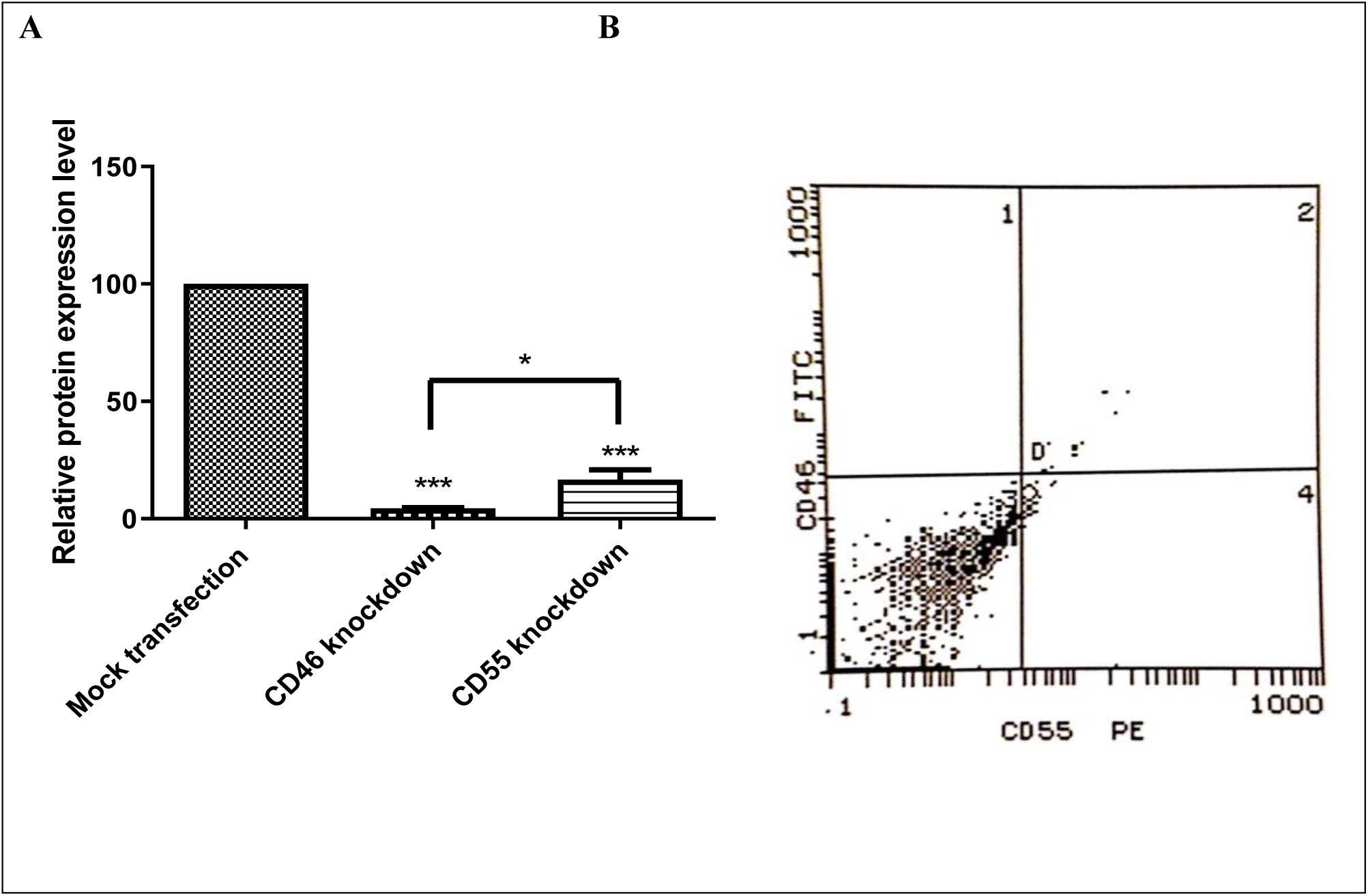
ShRNA-mediated combined knockdown of CD46 and CD55 mCRPs expression on HSB-2 cells compared to the mock transfected controls. (A) Co-silencing of CD46 and CD55 expression in transfected cells led to a highly significant reduction in the expression of both proteins(p<0.001). A significant difference was observed between CD46 and CD55 knockdown(p<0.05).The percentage of inhibition was calculated relative to nonsilencing shRNA controls (=100%). p<0.001(***), p<0.05(*). **(B)** Flow cytometry analysis chart for CD46 and CD55 expression on HSB-2 cells transfected with combined CD46 and CD55 shRNA silencing plasmids. Dots in the third quadrant show the double negative expression for both CD46 and CD55 after knockdown of both CD46 and CD55 genes using combined CD46 and CD55 silencing shRNA plasmids.

### Cell viability MTT assay following post transcriptional silencing of CD46 and CD55

In order to determine the effect of post transcriptional knockdown of CD46 and CD55 on cell viability in acute leukemia, MTT assay was performed where the absorbance of the formazan product was measured at 595 nm. Normal human serum (NHS) was used as a source of complement proteins. In the presence of NHS, a highly significant reduction in cell viability was observed in the CD46 silenced cells compared to untransfected (p<0.001) and mock transfected controls (p<0.05). Cell viability following post transcriptional silencing of CD46 was reduced by 71% compared to untransfected controls. Knockdown of CD55 resulted in a significant reduction of cell viability by 65%. Co-silencing of CD46 and CD55 has significantly reduced cell viability by 62% (Figure 8).

**Figure 8.**
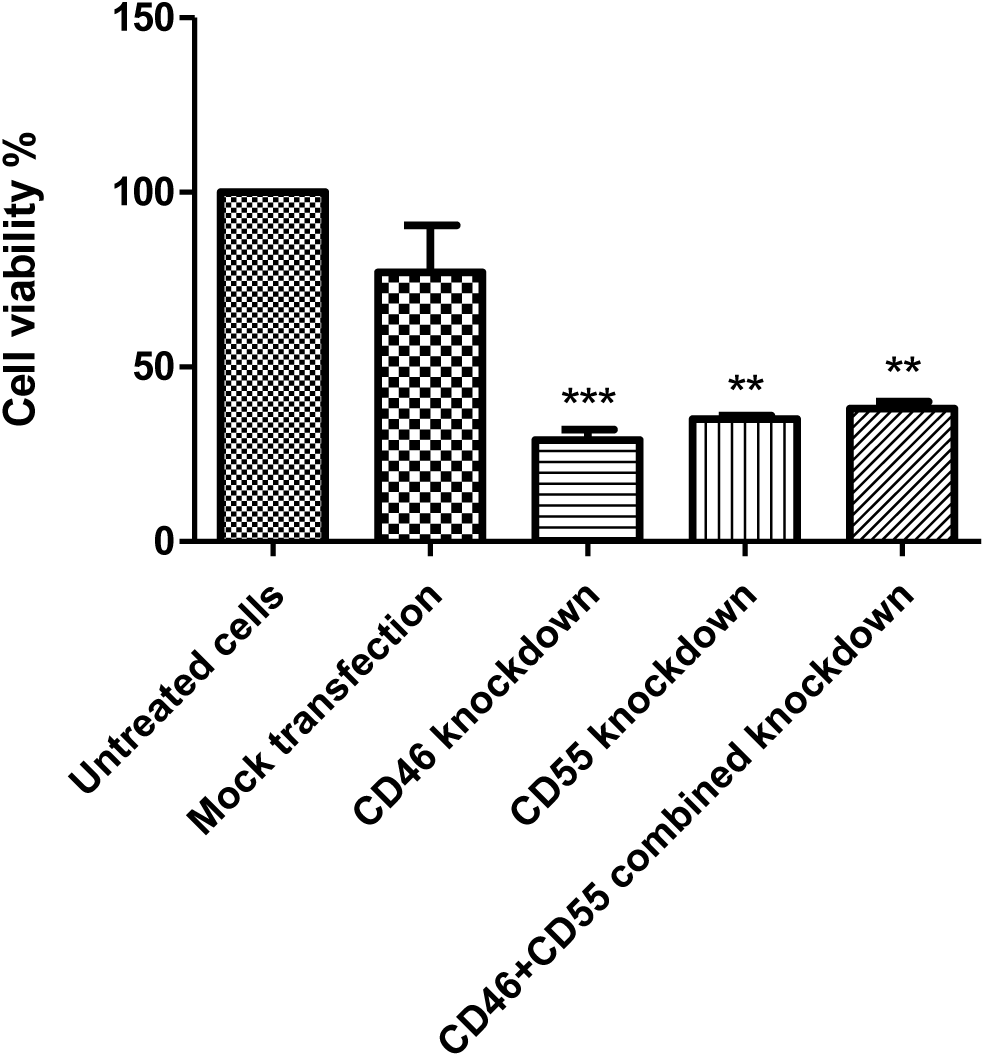
Acute lymphocytic leukemia cells viability after silencing of mCRPs. Viability of HSB-2 cells was analyzed after silencing of CD46 and CD55 protein expression separately as well as combined silencing of both proteins using shRNA plasmids. Cell viability was assessed using MTT reagent in the presence of NHS as a source of complement. Cells had a reduced viability in the absence of CD46 (p<0.001) and CD55 (p<0.01) as well as in CD46 and CD55 co-silenced cells (p<0.01). P<0.01 (**), p<0.001 (***).

## Discussion

The role of the complement system and the level of expression of its different regulatory proteins in various types of cancers has been an area of debate for many years. The complement components act as an important participant in the body’s immune surveillance against cancer. However, recent findings have suggested that the complement system can also have certain tumor-promoting capabilities ^[23]^.

Cancer cells have an increased capacity to activate the complement system, where some types of cancers such as breast cancer and papillary thyroid cancer were found to have deposits of C5b-9 MAC complex, C3 and C4 ^[24][25]^. It was also reported that the complement is activated in lung, digestive tract and brain tumors ^[26][27][28]^. These complement components are able to defend the body against cancer by complement-dependent cytotoxicity (CDC) and antibody-dependent cell-mediated cytotoxicity (ADCC) ^[29]^. Complement proteins can inhibit angiogenesis, where C5a stimulates monocytes to produce sVEGFR-1 which sequesters Vascular endothelial growth factor (VEGF) ^[30]^.

On the other hand, cancer cells use several strategies to resist complement attack. One of these strategies is the overexpression of mCRPs; CD46, CD55 and CD59 ^[31][32][33]^. This hinders the efficacy of cancer therapies that depend on the usage of monoclonal antibodies which activate the complement to perform CDC and ADCC and hence, reduce the cytotoxic effect against cancer cells ^[34]^.

Since there is an ever going debate as to whether the complement system is pro-or anti-cancer, we aimed in the current study to investigate the expression pattern of two important complement regulatory proteins; CD46 and CD55 in ALL and AML cancer patients.

To our knowledge, this is the first study of mCRPs in Egyptian cancer patients and one of very few studies that investigate their role in acute leukemia. qRT-PCR was carried out to determine the steady state levels of mRNA transcripts of both mCRPs in 30 acute leukemia patients from both genders in comparison to the expression pattern in healthy subjects. Our results showed a noticeable downregulation of CD46 mCRP in both acute leukemia types, ALL and AML compared to healthy controls where in AML samples CD46 expression was downregulated by 4 fold and in ALL it was downregulated by 5.5 fold compared to expression in healthy controls. The observed CD46 downregulation was more prominent among male patients compared to females. This observation suggests a hormonal link to the complement circuitry.

Our results agree with a previous study on B cell leukemia patients, also reporting CD46 downregulation. The expression level of CD46 was found to be relatively low on freshly isolated B cell leukemia samples compared to higher, but varying levels of CD55 and CD59 ^[35]^. In another study conducted on renal tumor cells, a low expression level of CD46 was associated with less advanced tumor growth ^[36]^. On the other hand, several earlier investigations on hematological malignancies reported that CD46 expression was 2-8 folds higher in ALL, AML, CLL and CML patients and in leukemia cell lines compared to normal cells ^[37][38]^. Another study conducted to investigate the level of mRNA expression of CRPs in ALL and AML patients compared to healthy subjects proved that there was no significant difference in any of the three groups examined for the expression level of CD46 ^[39]^. This shows that the expression of CD46 is rather heterogeneous in leukemic patients. One possible explanation for the variability in CD46 expression could be that no selective force is present to control the production of CD46 by cancerous cells. So apparently, without the presence of any external interference, the production of mCRPs will remain variable ^[32]^.

In the current study, the downregulation of CD46 is suggestive of a defense mechanism conducted by leukemic cells against ADCC (antibody dependent cell mediated cytotoxicity) where downregulation of CD46 could decrease the levels of iC3b molecules deposited on leukemic cells, which in turn can reduce their binding capabilities to their receptor, CR3 on effector immune cells, as natural killer cells. Consequently, the binding of monoclonal antibodies accumulating on the leukemic cell surface to their FCγ receptor on effector cells decreases, hence reducing the effect of ADCC ^[40][41]^.

CD55was also significantly down regulated (P<0.05) with approximately 3.8 fold in AML and ALL patients suggesting the possibility that cancer may evade complement attack mechanism and may even manipulate it to its benefit. Similar findings were reported in an earlier study where lymphoid cells especially non-Hodgkin’s lymphoma cells, were reported to lack CD55, although, in most cases, another phosphatidyl inositol-anchored protein, CD59, was still present. It was explained that levels of CD55 are not definitively regulated in tumor cell lines ^[37]^.

A significant reduction in the expression of both CD46 and CD55 proteins as well was confirmed by FACS analysis of peripheral blood samples of acute leukemia patients except for ALL patients where no significant difference was observed in CD55 protein expression between control groups and patients. These results further suggest that these mCRPs are mostly reduced in acute leukemia patients on both the transcriptome and proteome expression level.

The current results highlight the fact that the complement components can work in favor of cancerous cells inhibiting antitumor immunity as mentioned in previous studies, where the complement components levels are positively correlated with the size of a tumor as well as the progression of different types of cancers ^[42][43][44][45]^. C3a and C5a complement components have shown to act as key players supporting the development of cancer cells where C3aR/C5aR signaling on the surface of T lymphocytes acts as an immune checkpoint inhibiting the production of IL-10 in effector T lymphocytes ^[42]^. Although IL-10 is known for its immune suppressive function, it also acts to enhance activation and proliferation of certain immune cells in support of antitumor activity ^[46][47]^.Suppressing the production of IL-10 in T lymphocytes through C3aR/C5aR signaling greatly affects the antitumor activity and hence supports the growth of cancer cells ^[42]^ ^[48]^. Moreover, these findings are further supported by the fact that the co-stimulation of CD46 and T cell antigen receptor (TCR) on T cell surface induces the secretion of IL-10 ^[49]^. This is obviously reversed in the results of the current study due the remarkable reduction in CD46 expression level leading to a reduced level of IL-10 production and this means more growth of cancerous cells.

Gene specifics shRNA plasmids were also used in this study to knockdown the gene expression of two mCRPs; CD46 and CD55 in HSB-2, an ALL cell line. Expression level of both mCRPs was measured by flow cytometry following knockdown using CD46 and CD55 monoclonal antibodies to ensure successful transfection.

Cell viability was then evaluated in transfected cells using MTT assay in the presence of NHS as a source of complement proteins. Cell viability decreased by 71% in case of CD46 silencing and silencing of CD55 protein reduced cell viability by 65% (p<0.01) compared to untreated cells suggesting that CD46 and CD55 may be involved in protection of these cells against complement mediated attack at some point in time. Similarly in a study by N.Geis et al., small interfering RNAs (siRNAs) have been designed for posttranscriptional gene knock down of CD46, CD55 and CD59 targeting tumor cells’ sensitization to complement attack and thus to further exploit complement for tumor cell destruction. Upon mCRP knock down, complement dependent cytotoxicity (CDC) was augmented by up to 24±0.75% ^[50]^.

The contradicting roles of both mCRPs in acute leukemia may depend on the stage of cancer cells’ differentiation as well as the type of cancer itself ^[32]^. In addition to that, mCRPs expression differs according to host factors such as hormones and cytokines produced by neighboring cells ^[35][51]^. This raises the hypothesis that the complement system is likely to have a dual role in cancer.

In agreement with previous studies, our results proclaim dual roles of the complement system in cancer, where it confers a protective mechanism against tumorigenesis acting as a key player in tumor immunoediting through CDC and ADCC^[34][41]^. However, it also has a pro-tumorigenic potential in certain cancer stages and under certain conditions, from being an essential part in the inflammatory response which is important for tumor formation and progression to participating in fundamental hallmarks in cancer progression such as angiogenesis and metastasis ^[52][53][54]^. Furthermore, comparing our results with previous studies on the complement and its relation to different types of cancer shows how the responses of the complement components are unique in each type of cancer and how cancers can evolve to develop mechanisms that subvert the complement system to their benefit^[55]^.

This study highlights the role of complement regulatory proteins in the pathogenesis of cancer and its duality in the tumor microenvironment where a delicate balance between antitumor and tumor-promoting complement activities exists. In conclusion, the multifunctional properties of the complement system suggest that it has opposing roles in cancer where its biological functions are much more diverse than a simple elimination of target cells.

## Abbreviations

mCRPs: membrane complement regulatory proteins
RCA: regulators of complement activation
ALL: acute lymphocytic leukemia
AML: acute myelogenous leukemia
MCP: membrane co-factor protein
DAF: decay accelerating factor
GPI: glycosylphosphatidylinositol
NCI: national cancer institute
qRT-PCR: quantitative Real-time Polymerase chain reaction
GOI: gene of interest
HK: housekeeping gene
shRNA: short hairpin RNA
PEI: Polyethylenimine
FITC: fluorescein isothiocyanate
PE: phycoerythrin
NHS: normal human serum
CDC: complement-dependent cytotoxicity
ADCC: antibody-dependent cell-mediated cytotoxicity
VEGF: Vascular endothelial growth factor
CLL: chronic lymphocytic leukemia
CML: chronic myelogenous leukemia.

